# Single-nuclei sequencing of *Moricandia arvensis* reveals bundle sheath cell function in the photorespiratory shuttle of C_3_-C_4_ intermediate Brassicaceae

**DOI:** 10.1101/2024.12.02.626447

**Authors:** Sebastian Triesch, Vanessa Reichel-Deland, José Miguel Valderrama Martín, Michael Melzer, Urte Schlüter, Andreas P.M. Weber

## Abstract

Spatially confined gene expression determines cell identity and is fundamental to complex plant traits. In the evolutionary transition from C_3_ to the more efficient C_4_ photosynthesis, restricting the glycine decarboxylase reaction to bundle sheath cells initiates a carbon concentrating mechanism via the photorespiratory glycine shuttle. This evolutionary step is generally thought to play an essential role in the progression from ancestral C_3_ to C_4_ photosynthesis. Plants operating this shuttle are often referred to as C_3_-C_4_ intermediates or C_2_ species. Within the Brassicaceae family, which includes model plants and crops, such species have evolved independently at least five times. However, research on the biochemistry of C_3_-C_4_ intermediates in the Brassicaceae has been limited to a few case studies of differentially localized proteins between mesophyll and bundle sheath cells. Here, we leveraged recent advances in single-cell transcriptome sequencing to better understand how new cellular specialization affects interconnected pathways. We generated a single-nuclei RNA sequencing dataset for *Moricandia arvensis*, a Brassicaceae with C_3_-C_4_ intermediate characteristics, and compared it to a publicly available single-cell transcriptome of leaf tissue of the C_3_ *Arabidopsis thaliana*. We independently confirmed the localization of selected photorespiratory proteins by electron microscopy of immunogold-labelled leaf sections. Our analysis revealed the shift in expression of genes directly associated with the photorespiratory glycine decarboxylase reaction, but also of related pathways, such as ammonium assimilation, synthesis of specific amino acids, redox regulation, and transport to the *M. arvensis* bundle sheath. In contrast, the expression of these genes was not restricted to this cell type in the C_3_ plant.

**Highlight:** Single-nucleus RNA sequencing of *Moricandia arvensis* reveals bundle sheath cell-specific expression of photorespiration genes and associated pathways beyond glycine decarboxylase, including ammonium assimilation and redox regulation. This highlights the key role of metabolic compartmentalization in supporting C_3_-C_4_ intermediate photosynthesis.

## Introduction

Complex plant traits require distinct spatial gene regulation and C_3_-C_4_ intermediate photosynthesis is a highly distinctive trait that involves a specialized anatomy and metabolism (Giacomello, 2021; Lundgren, 2020; Schlüter and Weber, 2016). C_3_-C_4_ intermediate species, which have evolved multiple times in more than 50 species, are a valuable source for studying the early steps of C_4_ evolution and the genetics underlying cell-specific gene expression (Lundgren, 2020; Sage *et al*., 2018; Walsh *et al*., 2023). The Brassicaceae family contains at least five independent origins of C_3_-C_4_ intermediate photosynthesis, in phylogenetic proximity to well-characterized model species like *Arabidopsis thaliana* or *Arabis alpina,* and economically relevant crop species such as *Brassica oleracea* (cabbage), *Brassica napus* (rapeseed), or *Diplotaxis tenuifolia* (arugula) (Guerreiro *et al*., 2023). The close phylogenetic proximity of model-, crop- and C_3_-C_4_ plants makes the Brassicaceae family an ideal system to study the causes and consequences of differential cellular specialization.

In the Brassicaceae genus *Moricandia*, C_3_-C_4_ intermediate characteristics include a reduced carbon compensation point (Rawsthorne *et al*., 1988b) and an altered leaf ultrastructure, including bundle sheath cells (BSC) with a high number of centripetally arranged chloroplasts (Schlüter *et al*., 2017). It was also shown that the P-protein of the glycine decarboxylase complex (GDC) selectively localizes to the BSC mitochondria in C_3_-C_4_ intermediate *Moricandia*, but not in closely related C_3_ *Moricandia* species (Hylton *et al*., 1988; Rawsthorne *et al*., 1988a, b). GDC is a crucial component of photorespiration as it catalyzes the decarboxylation reaction of photorespiratory glycine to CO_2_, NH_3_ and 5,10-methyltetrahydrofolate which is converted to serine by a serine hydroxymethyltransferase (SHM). The BSC-specific activity of the GDC P-protein has been demonstrated in several C_3_- C_4_ intermediate and C_4_ species and it is widely agreed to be a critical step in the evolution of C_4_ photosynthesis via C_3_-C_4_ intermediate stages (Schulze *et al*., 2016). In the current model of the glycine shuttle, the absence of the GDC-P protein in the mesophyll cell (MC) of *M. arvensis* leads to the accumulation of glycine, which then passively diffuses into the BSC where it is selectively decarboxylated. This glycine shuttle and the associated BSC-specific release of CO_2_ leads to an elevated CO_2_ partial pressure around Rubisco and the photorespiratory CO_2_ can be efficiently refixed (Bauwe *et al*., 2010). The more efficient Rubisco carboxylation reaction is one of the potential advantages of the glycine shuttle in these C_3_-C_4_ intermediate species. This is accompanied by a high number and centripetal organization of organelles in the BSC that limits CO_2_ leakage.

Glycine decarboxylation also releases nitrogen in form of NH_3_, creating a nitrogen imbalance towards the BSC (Monson and Rawsthorne, 2000). Several routes for the back-shuttle of photorespiratory nitrogen have been proposed using flux modelling (Mallmann *et al*., 2014). They propose three scenarios for ammonium shuttles between the MC and the BSC: a glutamate/2-oxoglutarate shuttle, an alanine/pyruvate shuttle, and an aspartate/malate shuttle. Since aspartate, malate, pyruvate and alanine itself are exchanged between MC and BSC in C_4_ photosynthesis, the proposed nitrogen back-shuttles in C_3_-C_4_ intermediate plants could have primed primordial C_4_ cycles and facilitated the evolution of the additional anatomical and metabolic C_4_ traits. However, there is limited experimental evidence for the presence of these shuttles in *Moricandia,* other than increased metabolite levels of glutamate, malate and alanine, suggesting the operation of multiple or mixed shuttles in C_3_-C_4_ intermediate *M. arvensis* and *M. suffruticosa* (Schlüter *et al*., 2017).

The establishment of a glycine shuttle that initiates further steps of C_4_ evolution has been predicted by modelling studies, suggesting a smooth pathway to C_4_ (Heckmann *et al*., 2013). Previous conceptual models place anatomical preconditioning events, such as the increase of BSC organelles and their localization to the inner BSC, before the shift of GDC activity to the BSC (Sage, 2016). Not much is known about the genetic regulation underpinning the evolution of C_4_ anatomy. In C_4_ grasses such as maize and *Setaria*, the contribution of GOLDEN2-like and SCARECROW transcription factors (TFs) has been shown to influence BSC anatomy and chloroplast biogenesis (Lambret-Frotte *et al*., 2024; Slewinski *et al*., 2012) but the target genes controlled by these TFs and their cell-specific regulation remain unknown. The significant influence of TFs on the complex development of C_4_ anatomy indicates that multiple genes are affected by the evolution of C_4_ anatomy and that this evolution is likely mediated by changes in *cis*-regulatory factors (Swift *et al*., 2024).

We reasoned that to understand the genetic basis of C_3_-C_4_ intermediacy, we need to elucidate gene expression dynamics in a spatial context, in particular the shift in expression between MC and BSC. Methods for single-cell transcriptome studies in C_3_-C_4_ intermediate or C_4_ species were so far limited by the physical separation of BSC from phloem tissue or even BSC from MC (Aubry *et al*., 2014; Borba *et al*., 2023; Burgess *et al*., 2019). To date, laser microdissection or mechanical separation of bundle sheath strands have been used to isolate MC and BSC in C_4_ plants. These methods have provided valuable insights into the C_4_-specific transcriptional control at the single-cell level (Hua *et al*., 2021; Liu *et al*., 2022; Moreno-Villena *et al*., 2022). Isolation of tagged nuclei from specific cell types (INTACT) allows cell-type specific transcriptome sequencing, but requires the generation of transgenic material, which is not feasible for most plant species (Deal *et al*., 2020). Recent advances in the development of droplet-based single-cell or single-nuclei RNA sequencing (sc/snRNA-seq) methods have opened the door to the discovery of cell-specific transcriptome patterns at unprecedented resolution (Giacomello, 2021). snRNA-seq has recently been applied to C_4_ sorghum, providing deep insights into the evolution of differential partitioning of gene expression to the BSC from C_3_ rice (Swift *et al*., 2024). Applied to C_3_-C_4_ intermediate photosynthesis, snRNA-seq can answer important questions such as the interconnectivity of the glycine shuttle with other pathways and the consequences of photorespiratory specialization for the MC and BSC metabolism.

To address this, we performed snRNA-seq on leaf tissue from the C_3_-C_4_ intermediate plant *M. arvensis*. We established a nuclei isolation protocol for *M. arvensis* and utilized droplet-based snRNA-seq employing the 10X workflow. The resulting dataset allowed us to distinguish specific cell types and their corresponding transcriptional profiles. To confirm that the patterns we found were associated with C_3_-C_4_ intermediate traits, we compared the dataset with a publicly available *A. thaliana* leaf scRNA-seq dataset (Kim *et al*., 2021).

We examined the datasets with respect to three main research questions: (1) What is the role of the BSC in a leaf with C_3_-C_4_ intermediate photosynthesis compared to a C_3_ BSC? (2) How is the glycine shuttle integrated into the BSC-specific nitrogen metabolism in *M. arvensis*? (3) Can we infer *cis*-regulatory patterns underlying BSC-specific gene expression?

We determined the expression of genes involved in photorespiration and the proposed pathways for the transport of photorespiratory nitrogen. Comparison of the single-cell transcriptomes of *M. arvensis* and *A. thaliana* also revealed a remarkable recruitment of gene expression to the BSC from *A. thaliana* to *M. arvensis*. We found that this BSC functionalization was largely driven by a shift of gene expression associated with photorespiration, ammonium assimilation, redox balance, and transport to the *M. arvensis* BSC. It is generally accepted that the evolution of C_4_ photosynthesis via C_3_-C_4_ intermediate stages involves the recruitment of existing C_3_ genes into modified gene regulatory networks, probably *via* the exaptation of *cis*-elements (Hibberd and Covshoff, 2010). To this end, we analyzed *M. arvensis* upstream DNA sequences for the presence of motifs for spatially co-expressed TFs. Although limited by sequencing depth, we found the BSC-specific expression of the TGA and NAC TFs, which may mediate BSC specificity in C_3_-C_4_ intermediate *M. arvensis*.

## Material and Methods

### Plant growth

*M. arvensis* MOR1 seeds were sterilized using chlorine gas sterilization for 1 h and germinated on ½ strength Murashige and Skoog medium with 0.4 % (w/v) agar for 7 days at room temperature and a 12 h light-dark cycle. Germinated seedlings were transferred to soil and grown at a 12 h light-dark cycle and 25 °C (light) and 22 °C (dark).

### Nuclei extraction

2 g leaf material of the combined fifth and sixth leaf (counting from the cotyledon) were cut using scissors and placed on a petri dish containing 1 mL LB01 buffer (15 mM Tris-HCl pH = 7.5, 2 mM EDTA, 80 mM KCl, 20 mM NaCl, 15 mM 2-mercaptoethanol, 0.2 % (v/v) Triton X-100, 0.5 mM Spermine). Leaves were chopped on ice for 2 min with a razor blade. After addition of 1 mL LB01 buffer, leaves were chopped for another 3 min. 3 ml LB01 buffer were added to the chopped leaves in the petri dish and the mixture was incubated on ice for 15 min with gentle agitation every 3 min. The mixture was filtered through a 100 µm filter and subsequently through a 20 µm filter. All filters were pre-wetted with 1 ml LB01 buffer. The filtrate was overlayed on a density gradient centrifugation buffer (DGCB; 1.7 M sucrose, 10 mM Tris-HCl pH = 8.0, 2 mM MgCl_2_, 5 mM 2-mercaptoethanol, 1 mM EDTA, 0.15 % Triton X-100) and centrifuged at 4 °C and 1500 g for 30 min. The supernatant was discarded and the pellet was resuspended with 400 µl 10X nuclei resuspension buffer. A 30 µl aliquot of the nuclei suspension was stained with 1.5 µl DAPI to check nuclei integrity under a fluorescence microscope. All buffers were supplemented with 1 U/µl RNase inhibitor (Roche).

### Single Cell Library Generation

A total of 5,000 nuclei were used as input for the single-cell droplet library generation on the 10X Chromium Controller system utilizing the Chromium Single Cell 3’ NextGEM Reagent Kit v3.1 according to manufacturer’s instructions. Sequencing was carried out on a NextSeq 2000 system (Illumina Inc. San Diego, USA).

### Genome annotation and orthology analysis

The *A. thaliana* TAIR10 genome and annotation was obtained from www.arabidopsis.org (Berardini *et al*., 2015; Lamesch *et al*., 2012). The *M. arvensis* MOR1 genome was obtained from Guerreiro *et al*. (2023), the *de novo* annotation using *Helixer* (Holst *et al*., 2023; Stiehler *et al*., 2020) was obtained from Triesch *et al*. (2024). *gffread 0.12.7* was used to convert gff3 to gtf files to map snRNA-seq reads. Homologs between *A. thaliana* and *M. arvensis* were identified using *MMseqs2* (Steinegger and Söding, 2017). If multiple *M. arvensis* homologs for one *A. thaliana* gene were found, all were retained.

### Processing of 10X Genomics single cell data

Raw reads from the *M. arvensis* snRNA-seq experiment were processed using *Cell Ranger 6.1.2* (10X Genomics, CA, USA) using the ARC-v1 chemistry option. The count matrix for the *A. thaliana* scRNA-seq experiment was kindly provided by Dr. Ji-Yun Kim (HHU Düsseldorf, Kim *et al*. (2021)). RNA velocity was determined using *Velocyto 0.17.17* (La Manno *et al*., 2018) and *scvelo 0.3.1*. Further processing of the count matrix from *Cell Ranger* was performed using *scanpy 1.9.3*. Genes with annotated chloroplast and mitochondrial origin were removed. Genes expressed in fewer than 10 cells and cells with fewer than 200 genes expressed were removed. Cells were clustered using a graph-based approach using 50 principal components and a resolution of 0.25 for *M. arvensis* and 0.5 for *A. thaliana*.

### Cluster annotation and marker gene inference

Single-cell markers genes for *A. thaliana* were obtained from PlantscRNAdb (Chen *et al*., 2021) and the *M. arvensis* homologs for these genes were used for the respective datasets. In both datasets, within each cluster, marker genes were defined as genes with a *p* < 1e^-20^ and a positive log2-fold change. Each marker gene was tagged with one or more cell types from the PlantscRNAdb marker gene annotation. Fisher’s exact test was employed to find cell type annotation enrichments within the marker genes for the individual clusters. The enriched cell type annotation with the lowest p-value was naïvely selected for annotation of the respective cluster.

### Cross-species cell-type comparison

Marker genes for each cluster/cell-type were defined as genes with *p* < 1e^-20^ and a positive log2-fold change. Differentially expressed genes were determined from expression matrices using Mann-Whitney-U-Tests with a significance threshold of *p* < 1e^-20^. For the between-species analysis, gene expression as read counts per transcript was normalized to the highest expressed marker gene of the respective cluster. *MapMan* bin classifications for enrichment analyses were taken from Triesch *et al*. (2024). *MapMan* bin enrichment was performed using Fisher’s exact test, calculating the enrichment of target gene sets within the complete set of genes in the TAIR10 or *M. arvensis* annotation, respectively. Sankey plots were created using www.sankeymatic.com.

### *Cis*-element detection and cross-species analysis

Transcription factor binding sites for all upstream sequences were predicted using *FIMO* (Grant *et al*., 2011) from the *MEME suite 5.5.5* (Bailey *et al*., 2009). To this end, the 3000 bp upstream sequences for all annotated *M*. *arvensis* and *A. thaliana* genes were dumped using a custom *python* script. Motif clustering was performed on transcription factor motifs using the JASPAR database (Castro-Mondragon *et al*., 2022). To account for sequence composition a background model was generated using the *fasta-get-markov* tool from the *MEME suite 5.5.5*. The target upstream sequence, the background model and the database were used as input for *FIMO*.

### Immunogold labelling analysis of leaf tissue by light- and transmission electron microscopy

The central part of the 5^th^ leaf from *M. arvensis* and rosette leaves from *A. thaliana* of at least three independent plants were cut, chopped in 2 x 2 mm pieces and placed in a 2 ml tube containing the freshly prepared fixative solution (2% formaldehyde + 0.5 % glutaraldehyde in 0,05 M Na-cacodylate buffer, pH 7.2). Gentle vacuum infiltration of the fixative using a vacuum pump was followed by three rounds of microwave assisted fixation with a *Pelco 34700 Biowave Laboratory Microwave Processor* at 150 W for 1 min with 1 min rest between rounds. Afterwards, the samples were washed once with 0.05 M Na-cacodylate buffer and 3 times with distilled water, applying 150 W for 1 minute in each washing step. Samples were then dehydrated using an ethanol series of 30, 40, 50, 60, 70, 80, 90 and 100 %. In each dehydration step, 150 W were applied for 1 min followed by 15 min incubation on a shaker at room temperature. 100 % ethanol was substituted as many times as required in order to remove as much chlorophyll as possible. For ultrastructural analysis, sections were embedded in Spurr resin, sectioned and analysed as described by Kavka *et al*. (2024).

Immunocytochemistry sectioning and immunogold labelling was performed as described in Schwarz *et al*. (2015). Resin infiltration samples were incubated in immediately prepared 25, 50, 75 and 100% Lowicryl HM20 solution in ethanol. Samples were incubated overnight in 25 % HM20, followed by 4 hours in 50, 75 % and overnight incubation at 100 % HM20 before polymerization in fill gelatine capsules at -10 °C under UV light in a *Leica EM ASF2* (automated freeze substitution system). The excessive resin was trimmed creating a trapezoid surface using a *Leica EM Trim* trimming device. Sections of 0.70 µM thickness were obtained with the *Ultracut UCT ultramicrotome* using a *Diatome Ultra Knife*. Antibodies were obtained from Peter Westhoff (Heinrich Heine University Düsseldorf) and from Stefan Timm (University Rostock) in case of the serine hydroxymethyltransferase (SHM) antibody. They were used in a 1:50 dilution in antibody buffer and are listed in Supp. Table S1.

Counting of gold particles and the area of the organelles were determined using ImageJ (Schneider *et al*., 2012). The density of gold particles was calculated by counting the gold particles on the images at a magnification of 89.000 x and then calculating the number of particles per μm^2^. The gold particle density was measured on three different samples and a minimum of 45 BSC and MSC from the surrounding of a minimum of 9 different veins. The labelling density was calculated as a mean of number of gold particles per µm^2^ from at least 13 different organelles.

Statistical analyses for the gold particle count were performed using PRISM 8 (Graphpad, https://www.graphpad.com). Differences in labelling densities of the components of the GDC (GLDP1, GLDH, GLDT), SHM and hydroxypyruvate reductase (HPR) between MC and BSC were analyzed using student’s t test. Differences were considered to be statistically significant when the P value was < 0.01.

## Results

### Single-nuclei sequencing of *Moricandia arvensis*

To gain an understanding of the differential gene expression patterns in a leaf of a C_3_-C_4_ intermediate Brassicaceae species, the fifth and sixth leaves of young *M. arvensis* plants (counting from the first cotyledon) were used for nuclei extraction and snRNA-seq. Across three biological replicates, a total of 11,013 nuclei were used for snRNA-seq using the 10X single-nuclei workflow with a median of 1,070 identified genes per nucleus. The output was visualized using uniform manifold approximation projection (UMAP) and subjected to unsupervised clustering using *scanpy*, resulting in 10 distinct cell clusters (Fig. 1A, Supp. Material S1). The expression levels of the closest *M. arvensis* homologs for single-cell marker genes from *A. thaliana* (Chen *et al*., 2021) were used for annotation of cell types (Fig. 1B).

**Figure 1:**
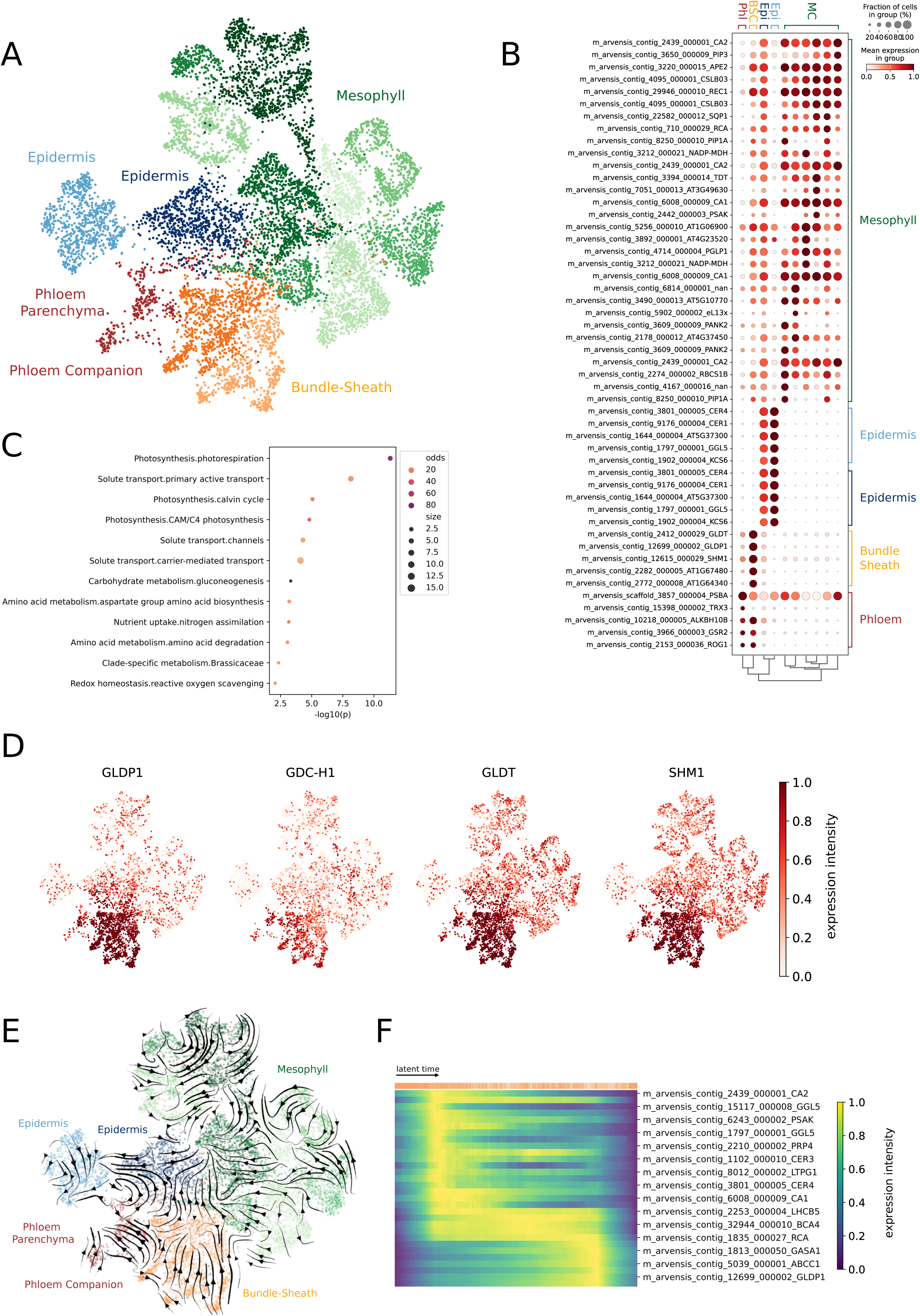
**A:** UMAP of transcript profiles for *Moricandia arvensis* 11,013 leaf cells. Cells were grouped into ten clusters using *scanpy*. Cell identities were assigned using marker genes for different cell types (**B**). **C:** Enrichment of MapMan bins for differentially expressed genes between *M. arvensis* bundle sheath and mesophyll clusters. **D:** UMAP with heatmap overlays indicating expression strength of selected genes. **E:** UMAP with RNA velocity arrows modelled using *velocyto* and *scvelo*. **F:** Heatmap of expression intensity for selected genes across a modelled latent time gradient for 1,687 bundle sheath cells.

In the resulting dataset, six MC clusters, one BSC cluster, one phloem and two epidermis cell clusters were identified. The MC clusters were identified by the strong expression of genes encoding CARBONIC ANHYDRASE homologs, subunits of photosystem 1, and RUBISCO ACTIVASE. The emergence of multiple MC subclusters could be attributed to the utilization of different biological replicates. Typically, the three biological replicates clustered together across all clusters, except for the mesophyll clusters, where the replicates showed more distinct clustering patterns (Supp. Fig. S1). However, the six mesophyll clusters showed the same expression patterns for most genes. Phloem clusters were identified by the expression of marker genes such as *TRX3*, *GSR2* and *ROG1*. The clusters were annotated as phloem parenchyma and phloem companion cells. Two distinct epidermis clusters were identified by expression of epidermal marker genes such as *ECERIFERUM* and *KCS6* (Fig. 1B). RNA velocity analysis indicated that the two clusters represented old and young epidermal cells rather than upper and lower epidermal layers (Fig. 1E).

1,687 cells formed a cluster that could be annotated as BSC tissue. Marker genes for the BSC cluster were the sub-proteins of the GDC and the *SULTR* sulfate transporter homologs. Sub-clustering of the BSC cluster using a higher clustering resolution resulted in two distinct BSC cell types. Latent time analysis using RNA velocity modeling showed that BSC from the two subclusters formed a linear gradient, suggesting that the two clusters did not originate from different developmental stages, but rather from different BSC types (Fig. 1F).

In order to determine the underlying genes for traits associated with C_3_-C_4_ intermediate photosynthesis, we calculated the differential expression of genes between MC and BSC clusters using Mann-Whitney U-tests with a significance threshold of *p* < 1e^-20^. We found that 219 genes were differentially expressed between MC and BSC, with 159 genes upregulated in BSC and 60 genes upregulated in MC. Enrichment analysis of the 219 differentially regulated genes revealed that within these genes the *MapMan* bins “Photosynthesis” and “Solute transport” were strongly enriched. The enrichment of genes within the “Solute transport” bin was mostly due to genes encoding the mitochondrial A BOUT DE SOUFFLE (BOU) transporter, the sulfate transporters SULTR2;2 and SULTR3;3, proton pump ATPases and transporters of the plasma membrane intrinsic protein (PIP) transporter family. The enrichment in the bin “Photosynthesis.photorespiration” was mainly due to the partitioning of different components of the GDC to the BSC. These included the genes encoding the GDC P-, H-, and T-protein (GLDP1, GDC-H1, GLDT) and the gene encoding the SHM (Fig. 1D). Using RNA velocity analysis, a pseudo-age was modelled for each cell. Within the BSC cluster the expression of *GLDP1* was observed in cells with a higher pseudo-age, whereas expression of the mesophyll marker gene *CARBONIC ANHYDRASE 2* (*CA2*) was observed in younger BSC (Fig. 1F).

### Comparative single-cell transcriptomics with *Arabidopsis thaliana*

To gain insight into cell-specific expression patterns that correlate with, and potentially underlie, traits associated with C_3_-C_4_ intermediate photosynthesis, we compared our *M. arvensis* snRNA-seq dataset with a publicly available single-cell RNA-seq (scRNA-seq) dataset from the C_3_ Brassicaceae species *A. thaliana*. This dataset contained expression data for 5,230 vasculature-enriched leaf protoplasts from two biological replicates (Kim *et al*., 2021). With a median of 3,342, the number of genes per cell was approximately three times higher in the *A. thaliana* protoplast dataset than in the *M. arvensis* nuclei population. To compare the datasets of both species, we assigned *M. arvensis* homologs of *A. thaliana* genes using *MMseqs2* (Steinegger and Söding, 2017). In this approach, genes are clustered based on protein sequences and ortholog and paralog sequences are combined.

For 20,512 *A. thaliana* genes, one or more homologs were found in *M. arvensis*. A median of two *M. arvensis* gene copies were found for each *A. thaliana* gene.

Using the same workflow as for the *M. arvensis* dataset, we visualized the *A. thaliana* scRNA-seq data using UMAP and clustered it into seven distinct clusters. However, when we attempted to combine the *A. thaliana* and *M. arvensis* datasets into a single UMAP projection, the combined cells clustered solely based on their respective species, rather than by the cell type. For this reason, both datasets were kept and clustered separately and comparisons were made on a per-cluster basis. The clusters were annotated using the same set of marker genes that was used for the analysis of *M. arvensis*. In contrast to the *M. arvensis* dataset, we were able to identify senescent, initial and guard cells in the *A. thaliana* single-cell population (Fig. 2A). Furthermore, the average sequencing depth reached approximately 96,000 reads per cell in the *A. thaliana* dataset, whereas it was limited to approximately 27,000 reads per cell in our *M. arvensis* dataset.

**Figure 2:**
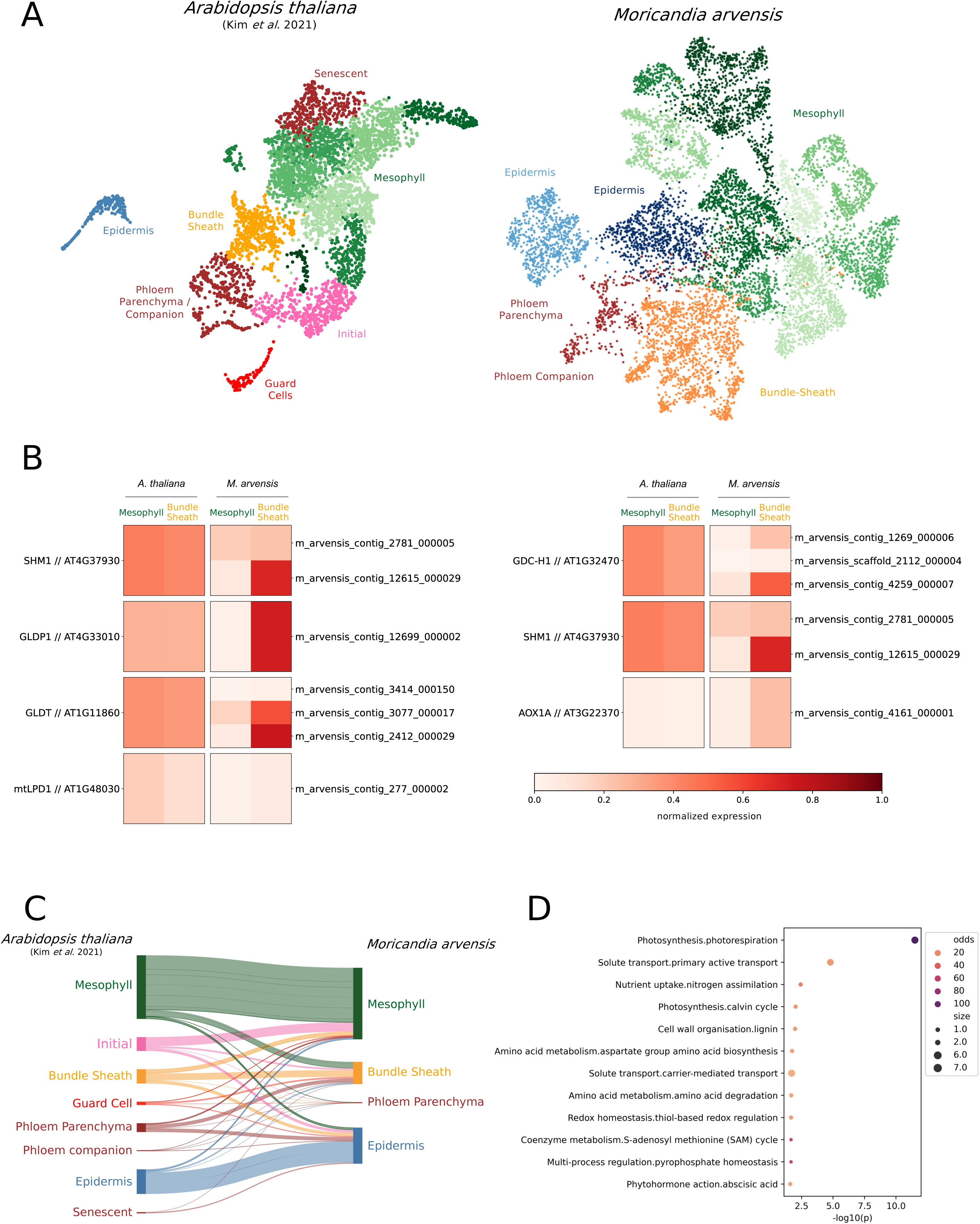
**A:** UMAP visualizations of the *Arabidopsis thaliana* single-cell RNA-seq dataset from Kim *et al*. (2021; left panel) and the *Moricandia arvensis* dataset from this study (right panel). **B:** Heatmaps indicating normalized expression in mesophyll and bundle sheath clusters in *A. thaliana* and *M. arvensis*. Expression was normalized to the highest expressed marker gene in the respective cluster and species. **C:** Sankey diagram indicating the expression of *M. arvensis* marker genes (right side of the diagram) in *A. thaliana* clusters (left side). **D:** Enrichment of *MapMan* bins for 101 differentially partitioned genes between *M. arvensis* and *A. thaliana* bundle sheath clusters.

Comparison of BSC populations between the species revealed that a proportion of *M. arvensis* BSC marker genes were not BSC-specific in *A. thaliana* (Fig. 2C). 71 *M. arvensis* BSC marker genes were also marker genes in the *A. thaliana* BSC and 101 *M. arvensis* BSC marker genes were not BSC-specific in *A. thaliana*. The 71 common BSC marker genes were enriched in the *MapMan* bins “Solute transport” and “Amino acid metabolism”, among others. We termed the 101 genes that showed cell-specific expression in only one of the species as “differentially partitioned”. These genes gained BSC-specific expression in *M. arvensis* during the evolutionary progression since the last common ancestor between the two plant species. The differentially partitioned *M. arvensis* BSC marker genes were enriched in the *MapMan* bins “Photosynthesis.photorespiration”, “Solute transport.Primary active transport” and “Nutrient uptake.nitrogen assimilation”. The enrichment of “Solute transport.Primary active transport” was largely due to the BSC-specific expression of ATP-binding cassette (ABC) transporters in *M. arvensis*. The enrichment of “Photosynthesis.photorespiration” in the set of differentially partitioned genes was due to homologs of the genes encoding subproteins of the GDC such as GLDP1, GLDT, and GDC-H1 (Fig. 2B). The BSC-preferential expression also diverged between *M. arvensis* homologs of single *A. thaliana* genes. For example, two copies of the *GLDT* gene were annotated in *M. arvensis*, one of which showed a stronger BSC-specific expression and one of which showed BSC-specificity with a stronger MC background. A similar pattern was found for the four *M.* arvensis paralogs of the single *A. thaliana NADP-MDH* gene, encoding the plastidial NADP-dependent malate dehydrogenase. Here, one paralog showed highly BSC-specific expression, whereas another paralog showed higher MC expression. Photorespiratory genes not directly involved in glycine decarboxylation were not differentially partitioned. Additionally, we found no evidence of increased *PHOSPHOENOLPYRUVATE CARBOXYLASE (PEPC)* transcripts in the MC that could indicate a C_4_-like shuttle (Supplemental Fig. S4).

### *Cis*-regulatory underpinnings of differential partitioning

Next, we examined the *cis*-regulatory landscape of differentially partitioned genes in the *M. arvensis* and *A. thaliana* datasets. To do this, the 3,000 bp upstream regions for each marker gene were analyzed using *FIMO* (Grant *et al*., 2011), which predicts TF binding sites. The list of putatively binding TFs based on upstream sequence elements was narrowed down using TF co-expression data from our single-cell RNAseq datasets.

For *A. thaliana*, binding sites for several co-expressed TFs were identified in the upstream regions of marker genes. A prime example was the accumulation of MYB30 TF binding sites in the *A. thaliana* epidermis marker genes, which was previously associated with epidermal wax synthesis (Zhang *et al*. (2019); Supp. Fig S6). Furthermore, DOF1 and DOF5 TFs were co-expressed and predicted to bind BSC marker genes in the *A. thaliana* data, which is supported by previous studies (Dai *et al*., 2022; Guo *et al*., 2009). The BSC-specific preference of DOF TFs was also found in a recent single-cell study of sorghum and rice (Swift *et al*., 2024). Recruitment of DOF TF binding sites is thought to have driven BSC repurposing in the evolution of C_4_ sorghum (Swift *et al*., 2024), for example by mediating differential partitioning of *NADP-ME* (Borba *et al*., 2023). A larger number of co-expressed and potentially binding TFs were identified in the *A. thaliana* dataset than in *M. arvensis*. This is likely an effect of the lower read coverage in the *M. arvensis* single-nuclei RNA-seq data, which is also likely the reason for the lack of detection of DOF TFs in the *M. arvensis* BSC.

Two co-expressed and putatively binding TFs were identified in the *M. arvensis* BSC marker genes, TGACG MOTIF-BINDING FACTOR 4 (TGA4) and NAC DOMAIN CONTAINING PROTEIN 83 (NAC083; Supp. Fig. S6). Interestingly, these TFs were not identified as co-expressed and binding in *A. thaliana* BSC marker genes.

### Leaf ultrastructure and protein localization through immunogold labelling

To corroborate the biological insights gained from our *M*. *arvensis* snRNA-seq dataset and the *A. thaliana* dataset from (Kim *et al*., 2021), we performed immunogold labelling to localize selected proteins predicted to be differentially distributed between the C_3_ and C_3_-C_4_ intermediate study systems.

In *M. arvensis*, BSC mitochondria were significantly labelled for GLDP, GLDH, GLDT and SHM (Figure 4), with significant differences from MC mitochondria (Figure 4, Table 1). For HPR, no significant differences in labelling were found between BSC and MC mitochondria for *M. arvensis* (Table 1). In *A. thaliana*, in contrast, no statistically significant differences were observed between the two cell types for the localization of GLDT, GLDH, SHM and HPR (Table 1).

**Figure 3:**
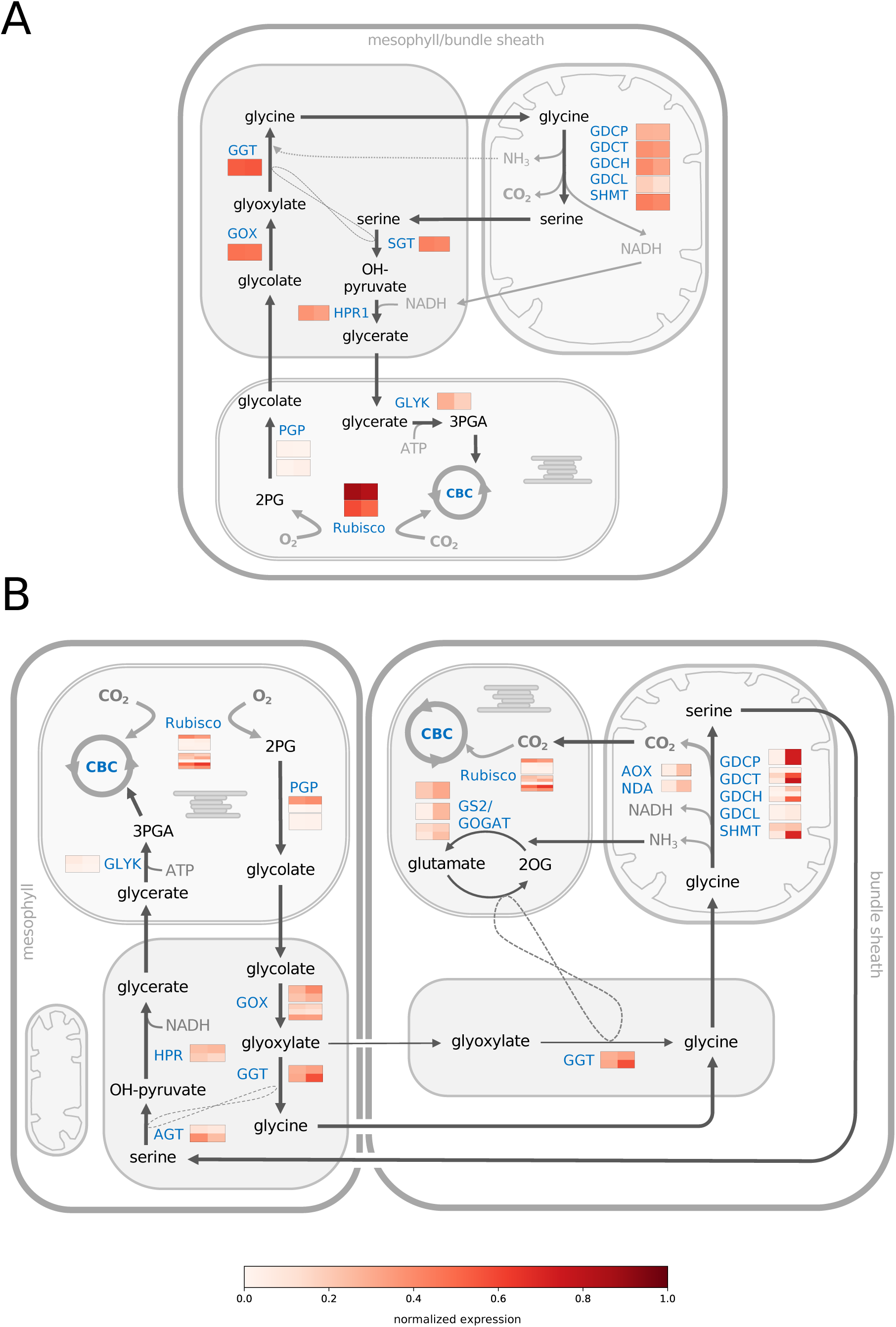
Schematic view of the photorespiratory pathway in *Arabidopsis thaliana* (**A**) and *Moricandia arvensis* (**B**) mesophyll and bundle sheath cells. Heatmaps indicate the expression of genes encoding selected photorespiratory enzymes from the respective *A. thaliana* and *M. arvensis* single-cell RNA-seq datasets. The expression was normalized to the highest marker genes in the respective cluster and species. The left tile of each heatmap indicates normalized expression in the mesophyll, the right tile indicates normalized expression in the bundle sheath. Multiple rows per tile indicate multiple gene copies. The nitrogen transfer reactions are shown in dashed lines. Abbreviations: CBC: Calvin-Benson-Bassham Cycle, 3-PGA: 3-Phosphoglyceric acid, Rubisco: Ribulose-1,5-bisphosphate carboxylase/oxygenase, GLYK: glycerate kinase, GOX: glycolate oxyidase, GGT: glutamate:glyoxylate aminotransferase, AGT: serine:glyoxylate aminotransferase, HPR: hydroxypyruvate reductase, GLDP/GLDT/GLDH/GLDL: glycine decarboxylase P/T/H/L protein, SHMT: serine hydroxymethyltransferase, PGLP: phosphoglycolate phosphatase, GOGAT: glutamine-2-oxoglutarate aminotransferase, GS: glutamine synthetase, 2OG: 2-oxoglutaric acid.

**Figure 4:**
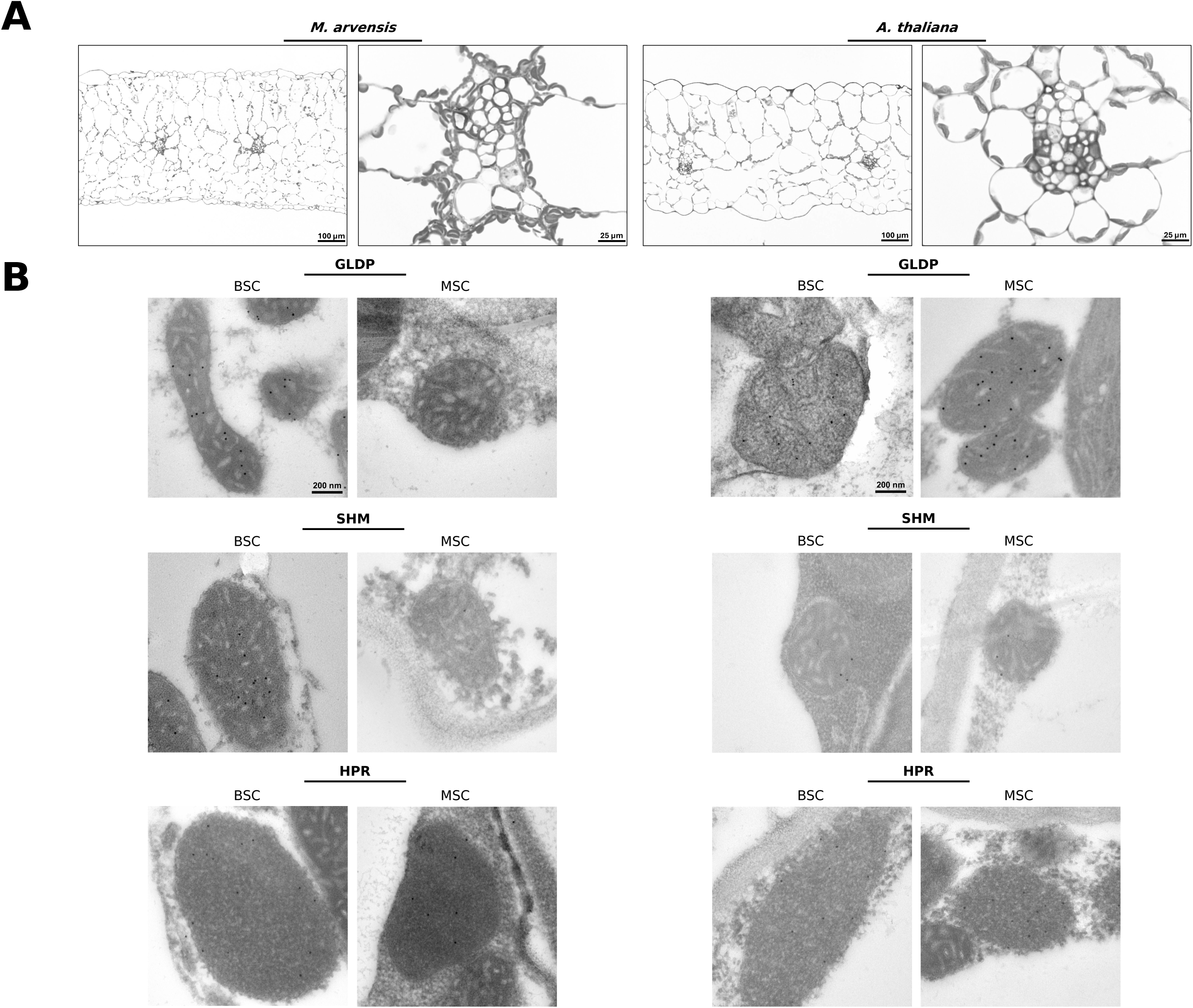
**A**: Light microscopy images of cross-sections of the central region of the fifth leaf from *Moricandia arvensis* and *Arabidopsis thaliana* embedded in SPURR resin. **B:** Electron microscopy images showing mitochondria of BSC and MC from *M. arvensis* and *A. thaliana*. Black dots are immunogold particles labelling selected photorespiratory proteins.

**Table 1:** Numbers of gold particles per unit area (µm^-2^) given as mean ± standard deviation. Data was obtained from different sections from at least three different leaves. Asterisks (*) indicate a significant difference (*P* < 0.01) between mesophyll (MC) and bundle-sheath cells (BSC) from the same species after t-test analysis.

|  | *A. thaliana* |  | *M. arvensis* |  |
| --- | --- | --- | --- | --- |
| Enzyme | Number of gold particles $\mu\text{m}^{-2}$ | | | |
|  | BSC | MC | BSC | MC |
| GLDP | 65.63±26.55 | 60.31±30.01 | 68.86±32.73* | 5.34±8.12* |
| GLDT | 36.99±19.35 | 35.02±17.70 | 109.81±43.29* | 21.16±11.44* |
| GDLH | 9.54±7.68 | 17.97±12.93 | 28.74±11.00* | 6.9±6.63* |
| SHM | 15.38±11.65 | 19.79±12.95 | 61.59±27.88* | 13.71±9.51* |
| HPR | 67.73±32 | 78.51±26.89 | 34.94±10.37 | 49.98±18.82 |

## Discussion

C_3_-C_4_ intermediate plants recapture the CO_2_ released by photorespiration more efficiently by operating a photorespiratory shuttle, also called C_2_ photosynthesis (Lundgren, 2020; Sage *et al*., 2012; Schlüter and Weber, 2016). The concentration of photorespiratory CO_2_ release in the BSC creates conditions that favor the carboxylation reaction of the BSC Rubisco. C_2_ biochemistry is mainly based on the specific localization of the GDC P-protein in BSC mitochondria (Rawsthorne *et al*., 1988b; Sage *et al*., 2012). This differential localization is observed in all C_3_-C_4_ intermediate taxa characterized to date, demonstrating that the evolution of this specialized trait relies on similar mechanisms (Hylton *et al*., 1988; Schlüter and Weber, 2016). Apart from the shift in GLDP protein localization, not much is known about cellular metabolic adjustments in the different evolutionary lineages. For C_3_-C_4_ Brassicaceae such as *M. arvensis* it has been suggested that differential gene expression is mainly limited to BSC specificity of the GLDP protein (Morgan *et al*., 1993). By using snRNA-seq of young *M. arvensis* leaves, we here provide a single-cell resolution transcriptome for a C_3_-C_4_ intermediate Brassicaceae species. Our dataset covers the majority of leaf cell types and shows robustness between the biological replicates (Supp. Fig. S1). We assigned cell identities to clusters based on marker genes, which are consistent with previous single-cell transcriptomes of plant leaves (Berrío *et al*., 2022; Procko *et al*., 2022; Swift *et al*., 2024).

To compare the single-cell transcriptome data of a C_3_-C_4_ photosynthetic leaf to a C_3_ species, we used publicly available data from *A. thaliana* leaf protoplasts (Kim *et al*., 2021). The split between the Brassiceae and the Camelineae tribes, which contain *M. arvensis* and *A. thaliana*, respectively, dates back to 25 Mya (Walden *et al*., 2020). However, the high number of shared marker genes between the respective cell types allows intra-species comparisons. Using the homologs for a set of *A. thaliana* marker genes, we were able to assign cell identities to the *M. arvensis* clusters with high fidelity. As the *A. thaliana* single-cell expression dataset was derived from leaf protoplasts, not nuclei, it had a higher read count per cell compared to our *M. arvensis* dataset, resulting in the detection of more transcripts per cell. For the highly expressed photorespiratory genes, the low read count in the *M. arvensis* dataset was not an obstacle, evidenced by the high number of differentially partitioned genes detected.

Due to the prominent role of the BSC in the C_3_-C_4_ intermediate leaf, we analyzed differential gene expression patterns between BSC in MC in *M. arvensis* as well as between the *M. arvensis* and *A. thaliana* BSC clusters. In doing so, we noticed that multiple BSC marker genes in *M. arvensis* were not BSC-specific in *A. thaliana.* The majority of these differentially partitioned genes showed MC and phloem parenchyma expression in *A. thaliana* (Fig. 2C). The reprogramming of the BSC by recruiting gene expression from MC and phloem cells was also observed in a comparative single-cell transcriptomic study between sorghum and rice (Swift *et al*., 2024) and it is thought to be one of the crucial drivers of C_4_ evolution (Hibberd and Covshoff, 2010; Reeves *et al*., 2017; Singh *et al*., 2023).

To find the genes underlying cellular specification during C_3_-C_4_ evolution in *M. arvensis*, we analyzed the set of differentially partitioned genes between the two species in greater detail. First, we observed a strong enrichment of photorespiratory gene expression in the *M. arvensis* BSC compared to *A. thaliana* (Fig. 1C). This was most pronounced in the differential partitioning of GDC proteins to the BSC of the C_3_-C_4_ intermediate leaf (Fig. 1D), but also included genes encoding the T-, L- and H-protein of the GDC, and of SHM, working together in the mitochondrial enzyme system. For GLDP, GLDT, GLDH and SHM this observation could also be confirmed on protein level by immunolocalization. The simultaneous differential partitioning of multiple parts of the GDC/SHM system is plausible because it would generate a fitness disadvantage to produce highly abundant proteins for a nonfunctioning GDC complex in the BSC. Loss of the GDC/SHM activity in the mesophyll cells leads to the induction of a glycine shuttle and increase of CO_2_ concentration in BSC by the enhanced photorespiratory glycine decarboxylation. Besides the enhanced CO_2_ release, the shift of the whole GDC/SHM complex to the BSC causes enhanced production of NH_3_, methylene-THF, and NADH in this cell type and this requires adjustments in other pathways.

Consistent with this, we observed increased expression of genes involved in the GLUTAMINE SYNTHETASE/GLUTAMINE-2-OXOGLUTARATE AMINOTRANSFERASE (GS/GOGAT) cycle in the BSC thus avoiding the loss of photorespiratory nitrogen by high ammonium fixation capacity directly in the BSC. In one photorespiratory cycle of C_3_ species like *A. thaliana,* two molecules of glycine are formed from glyoxylate by activity of the GLUTAMATE:GLYOXYLATE AMINOTRANSFERASE (GGT). Amino groups for this reaction are normally supplied by the SERINE:GLYOXYLATE AMINOTRANSFERASE (AGT). However, since only one molecule of serine is formed from two molecules of glycine by the mitochondrial GDC/SHM, additional amino donors must be imported into the peroxisome and these are mainly provided by the activity of the GS/GOGAT cycle (Figure 3). Interestingly, our dataset also indicated a BSC preferential expression of one of the *GGT* gene copies in *M. arvensis* (Figure 3). Thus, under operation of the photorespiratory glycine - serine shuttle, serine returning to the MC could provide some of the amino groups, but some of the glyoxylate (and possibly also some glycolate) could be transported to the BSC where amino donors from ammonium refixation products would be available for GGT activity (Figure 3, Supp. Fig. S3, Supp. Fig. S4). Both peroxisomal amino transferases accept various substrates as amino donors, thereby increasing the robustness of the system (Bauwe, 2023). A partial shift of GGT activity to the BSC of C_2_ species would also significantly reduce the nitrogen imbalance between the two cell types.

Our hypothesis is consistent with our snRNA-seq data, where we find a BSC-preferential expression of *GLYCOLATE OXIDASE* (*GOX*), *GGT* and *GS/GOGAT*. These data are well supported by early protein activity studies where Rawsthorne *et al*. (1988a, 1988b) used a mixture of protoplasting and strand-separation techniques to distinguish MC from BSC cells. Higher enzyme activities were found in *M. arvensis* BSC fractions for GDC, GOGAT, GS, but not for HPR and GOX compared to the MC fractions.

However, not all photorespiratory intermediates are recycled and, in particular, glycine and serine are removed from the cycle (Busch, 2020; Walker *et al*., 2024). In the source leaf, the BSC would also supply metabolites to other plant parts via the adjacent vein (Supp. Figure S3) for which the system could be buffered by the operation of additional metabolite transport such as the proposed glutamate/2-oxoglutarate, alanine/pyruvate and aspartate/malate shuttles (Mallmann *et al*., 2014). The shuttles could operate at low levels and would not require additional cell specific adaptations of the associated enzymes ALANINE AMINOTRANSFERASE (AlaAT), ASPARTATE AMINOTRANSFERASE (AspAT) and MALATE DEHYDROGENASE (MDH) (Supp. Figure S3).

Based on work in *Flaveria*, the C_4_ evolutionary model of Mallmann *et al*. (2014) further suggests that a MC-specific increase in PEPC activity could contribute to the nitrogen shuttle and further increases in CO_2_ in the BSC through additional decarboxylating enzymes. However, in *M. arvensis*, no increase in PEPC activity was found in leaf extracts (Schlüter *et al*., 2017), and our snRNA-seq data also shows no cellular preferences of enzymes that would be required for operating a C_4_-like shuttle (Suppl Figure S3, S4).

The mitochondrial GDC/SHM complex also provides backbones for the C_1_ metabolism and the BSC specificity of methionine synthases in our dataset indicates that synthesis of specific amino acids is also adjusted to the spatial distribution of precursors in the C_3_-C_4_ intermediate leaf (Supp. Figure S3).

Interestingly, expression of the gene encoding the BOU transporter that has been linked to GDC activity is also more abundantly expressed in the BSC. Eisenhut *et al*. (2013) observed that *BOU* expression correlates with the expression of genes encoding GDC proteins and *bou* mutants exhibit decreased GDC activity. The authors suggested that BOU may transport a GDC cofactor into the mitochondria. It is conceivable that BOU transports glutamate for the polyglutamylation of THF, a co-substrate for the GDC and SHM. The differential distribution of BOU in the C_3_-C_4_ intermediate BSC may ensure BSC-specific GDC activity and may influence C_1_ metabolism.

Since the mitochondrial photorespiratory reactions are also connected to NADH formation, it was plausible to assume that changes in the BSC metabolism between the C_3_ *A. thaliana* and the C_3_-C_4_ intermediate *M. arvensis* require changes in redox balancing and energy homeostasis. Our dataset revealed that genes encoding proteins involved in alternative mitochondrial respiratory chains such as ALTERNATIVE OXIDASE 1 A (AOX1A) and ALTERNATIVE NAD(P)H DEHYDROGENASE 1 (NAD1) were differentially partitioned towards the *M. arvensis* BSC. An upregulation of *AOX* gene expression was already observed in bulk transcriptome studies in *M. arvensis* but not in the C_3_ sister species *M. moricandioides*, indicating enhanced re-balancing of the redox metabolism due to the glycine shuttle (Schlüter *et al*., 2017). Redox homeostasis can also be maintained by the malate valve, especially through the interplay of NADP-MDH (NADP-dependent malate dehydrogenase) enzymes (Selinski and Scheibe, 2019). Whereas the plastidial NADP-MDH is encoded by a single gene in *A. thaliana*, three paralogs exist in *M. arvensis*. Interestingly, we found a differential partitioning between these paralogs in *M. arvensis*. This could hint at a tight control of the malate metabolism between MC and BSC, carried out by multiple genes with potentially individual regulatory mechanisms.

Similar to the situation in the C_3_ species *A. thaliana*, we found a higher BSC expression of genes involved in sulfur metabolism in *M. arvensis* (Supp. Fig. S2). This included especially the sulfate transport steps mediated by SULTR2;2 and SULTR3;3 as well as primary reductive steps of sulfate assimilation that were highly BSC-specific in both species. This specificity, however, was more pronounced in *A. thaliana*.

We also interrogated pathways that showed differential activity in MC and BSC of C_4_ species such as the Calvin-Benson cycle or sucrose and starch synthesis (Schlüter and Weber, 2020), but none of these could be assigned to BSC our *M. arvensis* dataset.

Taken together, our data suggest that the photorespiratory shuttle is softly integrated into the cell-specific *M. arvensis* leaf biochemistry. Besides the core GDC/SHM enzyme system, shifts in cell specificity were mainly found for interacting pathways such as ammonium assimilation, redox and energy balancing, transport processes and C_1_ metabolism. Such economical metabolic integration of the photorespiratory shuttle could be linked to cell specific regulatory elements in the underlying genome (Swift *et al*., 2024), providing plants with the ability to avoid the production of superfluous proteins.

The lack of a strong C_4_-like N-balancing shuttle would explain the evolutionary stability of C_3_-C_4_ intermediate Brassicaceae with a presumably low selection for further C_4_ cycle adjustments. In comparison with the engineering of a C_4_ pathway into C_3_ leaves, the C_2_ cycle in *M. arvensis* requires fewer genetic adjustments and strengthens the idea that C_3_-C_4_ intermediate traits are promising targets for engineering into a C_3_ crop chassis (Ermakova *et al*., 2020).

## Author contributions

**S.T.** conducted nuclei isolation, data analysis and wrote the manuscript. **V.R.D.** established the nuclei isolation workflow and assisted in nuclei preparation. **J.M.V.M.** and **M.M.** performed immunogold labelling assays. **U.S.** contributed to writing the manuscript. **U.S.** and **A.P.M.W.** conceptualized the project and supervised the work.

## Acknowledgements

The authors thank Dr. Yi-Jun Kim for providing the *A. thaliana* scRNA-seq transcriptome, Samantha Flachbart for assisting in nuclei preparation, Jana Peter for assisting in plant handling and Marion Benecke and Kirsten Hoffie for assisting in immunogold labelling assays. Peter Westhoff (Universität Düsseldorf) and Stefan Timm (University Rostock) provided the specific antibodies for immunolocalization. The authors furthermore thank Dr. Tobias Lautwein and Anja Schuster for help and technical support with the 10X single-nuclei workflow. Computational infrastructure and support were provided by the Centre for Information and Media Technology at Heinrich Heine University Düsseldorf. This work was funded by the Deutsche Forschungsgemeinschaft (German Research Foundation) under Germany’s Excellence Strategy EXC-2084/1 (project ID 390686111) and under the CRC TRR 341 (project ID 456082119).

## Data availability

Sequencing data is available in the NCBI sequence read archive (SRA) under the project ID PRJNA1186371.

## Literature

Aubry S, Smith-Unna RD, Boursnell CM, Kopriva S, Hibberd JM. 2014. Transcript residency on ribosomes reveals a key role for the Arabidopsis thaliana bundle sheath in sulfur and glucosinolate metabolism. Plant Journal 78, 659–673.

Bailey TL, Boden M, Buske FA, Frith M, Grant CE, Clementi L, Ren J, Li WW, Noble WS. 2009. MEME Suite: Tools for motif discovery and searching. Nucleic Acids Research.

Bauwe H. 2023. Humboldt Review: Photorespiration – Rubisco’s repair crew. Journal of Plant Physiology 280, 153899.

Bauwe H, Hagemann M, Fernie AR. 2010. Photorespiration: players, partners and origin. Trends in Plant Science, Vol. 15: Elsevier Current Trends, 330-336.

Berardini TZ, Reiser L, Li D, Mezheritsky Y, Muller R, Strait E, Huala E. 2015. The arabidopsis information resource: Making and mining the “gold standard” annotated reference plant genome. genesis 53, 474–485.

Berrío RT, Verstaen K, Vandamme N, Pevernagie J, Achon I, van Duyse J, van Isterdael G, Saeys Y, de Veylder L, Inzé D, Dubois M. 2022. Single-cell transcriptomics sheds light on the identity and metabolism of developing leaf cells. Plant Physiology 188, 898–918.

Borba AR, Reyna-Llorens I, Dickinson PJ, Steed G, Gouveia P, Górska AM, Gomes C, Kromdijk J, Webb AAR, Saibo NJM, Hibberd JM. 2023. Compartmentation of photosynthesis gene expression in C4 maize depends on time of day. Plant Physiology 193, 2306–2320.

Burgess SJ, Reyna-Llorens I, Stevenson SR, Singh P, Jaeger K, Hibberd JM. 2019. Genome-Wide Transcription Factor Binding in Leaves from C3 and C4 Grasses. The Plant Cell 31, 2297–2314.

Busch FA. 2020. Photorespiration in the context of Rubisco biochemistry, CO(2) diffusion and metabolism. Plant J 101, 919-939.

Castro-Mondragon JA, Riudavets-Puig R, Rauluseviciute I, Berhanu Lemma R, Turchi L, Blanc-Mathieu R, Lucas J, Boddie P, Khan A, Perez NM, Fornes O, Leung TY, Aguirre A, Hammal F, Schmelter D, Baranasic D, Ballester B, Sandelin A, Lenhard B, Vandepoele K, Wasserman WW, Parcy F, Mathelier A. 2022. JASPAR 2022: the 9th release of the open-access database of transcription factor binding profiles. Nucleic Acids Research 50, D165–D173.

Chen H, Yin X, Guo L, Yao J, Ding Y, Xu X, Liu L, Zhu Q-H, Chu Q, Fan L. 2021. PlantscRNAdb: A database for plant single-cell RNA analysis.

Dai X, Tu X, Du B, Dong P, Sun S, Wang X, Sun J, Li G, Lu T, Zhong S, Li P. 2022. Chromatin and regulatory differentiation between bundle sheath and mesophyll cells in maize. Plant Journal 109, 675–692.

Deal RB, Henikoff S, Division S, Hutchinson F. 2020. The INTACT method for cell type-specific gene expression and chromatin profiling in Arabidopsis. 6, 56–68.

Eisenhut M, Planchais S, Cabassa C, Guivarc’H A, Justin AM, Taconnat L, Renou JP, Linka M, Gagneul D, Timm S, Bauwe H, Carol P, Weber APM. 2013. Arabidopsis A BOUT DE SOUFFLE is a putative mitochondrial transporter involved in photorespiratory metabolism and is required for meristem growth at ambient CO₂ levels. The Plant journal : for cell and molecular biology 73, 836–849.

Ermakova M, Danila FR, Furbank RT, von Caemmerer S. 2020. On the road to C4 rice: advances and perspectives. Plant Journal 101, 940–950.

Giacomello S. 2021. A new era for plant science: spatial single-cell transcriptomics. Current Opinion in Plant Biology 60, 102041–102041.

Grant CE, Bailey TL, Noble WS. 2011. FIMO: scanning for occurrences of a given motif. Bioinformatics 27, 1017–1018.

Guerreiro R, Bonthala VS, Schlüter U, Hoang NV, Triesch S, Schranz ME, Weber APM, Stich B. 2023. A genomic panel for studying C3-C4 intermediate photosynthesis in the Brassiceae tribe. Plant Cell and Environment 46, 3611–3627.

Guo Y, Qin G, Gu H, Qu LJ. 2009. Dof5.6/HCA2, a Dof Transcription Factor Gene, Regulates Interfascicular Cambium Formation and Vascular Tissue Development in Arabidopsis. The Plant Cell 21, 3518–3534.

Heckmann D, Schulze S, Denton A, Gowik U, Westhoff P, Weber APM, Lercher MJ. 2013. Predicting C4 photosynthesis evolution: Modular, individually adaptive steps on a mount fuji fitness landscape. Cell 153, 1579–1579.

Hibberd JM, Covshoff S. 2010. The regulation of gene expression required for C4 photosynthesis. Annual Review of Plant Biology 61, 181–207.

Holst F, Bolger A, Günther C, Maß J, Triesch S, Kindel F, Kiel N, Saadat N, Ebenhöh O, Usadel B, Schwacke R, Bolger M, Weber APM, Denton AK. 2023. Helixer–de novo Prediction of primary eukaryotic gene models conbining deep learning and a Hidden Marcov Model. bioRxiv.

Hua L, Stevenson SR, Reyna-Llorens I, Xiong H, Kopriva S, Hibberd JM. 2021. The bundle sheath of rice is conditioned to play an active role in water transport as well as sulfur assimilation and jasmonic acid synthesis. Plant Journal 107, 268–286.

Hylton CM, Rawsthorne S, Smith AM, Jones DA, Woolhouse HW. 1988. Glycine decarboxylase is confined to the bundle sheath cells of leaves of C3-C4 intermediate species. Planta 175, 452–459.

Kavka M, Balles A, Bohm C, Dehmer KJ, Fella C, Rose F, Saal B, Schulze S, Willner E, Melzer M. 2024. Phenotypic screening of seed retention and histological analysis of the abscission zone in Festuca pratensis and Lolium perenne. BMC Plant Biol 24, 577.

Kim JY, Symeonidi E, Pang TY, Denyer T, Weidauer D, Bezrutczyk M, Miras M, Zöllner N, Hartwig T, Wudick MM, Lercher M, Chen LQ, Timmermans MCP, Frommer WB. 2021. Distinct identities of leaf phloem cells revealed by single cell transcriptomics. Plant Cell 33, 511–530.

La Manno G, Soldatov R, Zeisel A, Braun E, Hochgerner H, Petukhov V, Lidschreiber K, Kastriti ME, Lönnerberg P, Furlan A, Fan J, Borm LE, Liu Z, van Bruggen D, Guo J, He X, Barker R, Sundström E, Castelo-Branco G, Cramer P, Adameyko I, Linnarsson S, Kharchenko PV. 2018. RNA velocity of single cells. Nature 2018 560:7719 560, 494-498.

Lambret-Frotte J, Smith G, Langdale JA. 2024. GOLDEN2-like1 is sufficient but not necessary for chloroplast biogenesis in mesophyll cells of C4 grasses. The Plant Journal 117, 416–431.

Lamesch P, Berardini TZ, Li D, Swarbreck D, Wilks C, Sasidharan R, Muller R, Dreher K, Alexander DL, Garcia-Hernandez M, Karthikeyan AS, Lee CH, Nelson WD, Ploetz L, Singh S, Wensel A, Huala E. 2012. The Arabidopsis Information Resource (TAIR): Improved gene annotation and new tools. Nucleic Acids Research 40.

Liu WY, Yu CP, Chang CK, Chen HJ, Li MY, Chen YH, Shiu SH, Ku MSB, Tu SL, Lu MYJ, Li WH. 2022. Regulators of early maize leaf development inferred from transcriptomes of laser capture microdissection (LCM)-isolated embryonic leaf cells. Proceedings of the National Academy of Sciences of the United States of America 119, e2208795119–e2208795119.

Lundgren MR. 2020. C2 photosynthesis: a promising route towards crop improvement? New Phytologist 228, 1734–1740.

Mallmann J, Heckmann D, Bräutigam A, Lercher MJ, Weber APM, Westhoff P, Gowik U. 2014. The role of photorespiration during the evolution of C4 photosynthesis in the genus Flaveria. eLife 2014, 1–23.

Monson RK, Rawsthorne S. 2000. CO2 Assimilation in C3-C4 Intermediate Plants. 533-550.

Moreno-Villena JJ, Zhou H, Gilman IS, Lori Tausta S, Maurice Cheung CY, Edwards EJ. 2022. Spatial resolution of an integrated C4+CAM photosynthetic metabolism. Science Advances 8, 2349–2349.

Morgan CL, Turner SR, Rawsthorne S. 1993. Coordination of the cell-specific distribution of the four subunits of glycine decarboxylase and of serine hydroxymethyltransferase in leaves of C3-C4 intermediate species from different genera. Planta 190, 468–473.

Procko C, Lee T, Borsuk A, Bargmann BOR, Dabi T, Nery JR, Estelle M, Baird L, O’Connor C, Brodersen C, Ecker JR, Chory J. 2022. Leaf cell-specific and single-cell transcriptional profiling reveals a role for the palisade layer in UV light protection. The Plant Cell 34, 3261–3279.

Rawsthorne S, Hylton CM, Smith AM, Woolhouse HW. 1988a. Distribution of photorespiratory enzymes between bundle sheath and mesophyll cells in leaves of the C3-C4 intermediate species Moricandia arvensis (L.) DC. Planta 176, 527–532.

Rawsthorne S, Hylton CM, Smith AM, Woolhouse HW. 1988b. Photorespiratory metabolism and immunogold localization of photorespiratory enzymes in leaves of C3 and C3-C4 intermediate species of Moricandia. Planta 173, 298–308.

Reeves G, Grangé-Guermente MJ, Hibberd JM. 2017. Regulatory gateways for cell-specific gene expression in C4 leaves with Kranz anatomy. Journal of Experimental Botany 68, 107–116.

Sage RF. 2016. A portrait of the C4 photosynthetic family on the 50th anniversary of its discovery: Species number, evolutionary lineages, and Hall of Fame. Journal of Experimental Botany 67, 4039–4056.

Sage RF, Monson RK, Ehleringer JR, Adachi S, Pearcy RW. 2018. Some like it hot: the physiological ecology of C4 plant evolution. Oecologia 2018 187:4 187, 941-966.

Sage RF, Sage TL, Kocacinar F. 2012. Photorespiration and the evolution of C4 photosynthesis. Annual Review of Plant Biology 63, 19–47.

Schlüter U, Bräutigam A, Gowik U, Melzer M, Christin PA, Kurz S, Mettler-Altmann T, Weber APM. 2017. Photosynthesis in C3-C4 intermediate Moricandia species. Journal of Experimental Botany 68, 191–206.

Schlüter U, Weber APM. 2016. The Road to C4 Photosynthesis: Evolution of a Complex Trait via Intermediary States. Plant and Cell Physiology 57, 881–889.

Schlüter U, Weber APM. 2020. Regulation and Evolution of C 4 Photosynthesis. 1-33.

Schneider CA, Rasband WS, Eliceiri KW. 2012. NIH Image to ImageJ: 25 years of image analysis. Nat Methods 9, 671–675.

Schulze S, Westhoff P, Gowik U. 2016. Glycine decarboxylase in C3, C4 and C3–C4 intermediate species. Current Opinion in Plant Biology 31, 29-35.

Schwarz N, Armbruster U, Iven T, Bruckle L, Melzer M, Feussner I, Jahns P. 2015. Tissue-specific accumulation and regulation of zeaxanthin epoxidase in Arabidopsis reflect the multiple functions of the enzyme in plastids. Plant Cell Physiol 56, 346–357.

Selinski J, Scheibe R. 2019. Malate valves: old shuttles with new perspectives. Plant Biology (Stuttgart, Germany) 21, 21–21.

Singh P, Stevenson SR, Dickinson PJ, Reyna-llorens I, Tripathi A, Reeves G, Schreier TB, Hibberd JM. 2023. C 4 gene induction during de-etiolation evolved through changes in cis to allow integration with ancestral C 3 gene regulatory networks. 9.

Slewinski TL, Anderson AA, Zhang C, Turgeon R. 2012. Scarecrow Plays a Role in Establishing Kranz Anatomy in Maize Leaves. Plant and Cell Physiology 53, 2030–2037.

Steinegger M, Söding J. 2017. MMseqs2 enables sensitive protein sequence searching for the analysis of massive data sets. Nature Biotechnology 2017 35:11 35, 1026-1028.

Stiehler F, Steinborn M, Scholz S, Dey D, Weber APM, Denton AK. 2020. Helixer: Cross-species gene annotation of large eukaryotic genomes using deep learning. Bioinformatics.

Swift J, Luginbuehl LH, Hua L, Schreier TB, Donald RM, Stanley S, Wang N, Lee TA, Nery JR, Ecker JR, Hibberd JM. 2024. Exaptation of ancestral cell-identity networks enables C(4) photosynthesis. Nature.

Triesch S, Denton AK, Bouvier JW, Buchmann JP, Reichel-Deland V, Guerreiro RNFM, Busch N, Schlüter U, Stich B, Kelly S, Weber APM. 2024. Transposable elements contribute to the establishment of the glycine shuttle in Brassicaceae species. Plant Biology 26, 270–281.

Walden N, German DA, Wolf EM, Kiefer M, Rigault P, Huang XC, Kiefer C, Schmickl R, Franzke A, Neuffer B, Mummenhoff K, Koch MA. 2020. Nested whole-genome duplications coincide with diversification and high morphological disparity in Brassicaceae. Nature Communications 11.

Walker B, Schmiege SC, Sharkey TD. 2024. Re-evaluating the energy balance of the many routes of carbon flow through and from photorespiration. Plant Cell Environ 47, 3365–3374.

Walsh CA, Bra, die, utigam A, Roberts MR, Lundgren MR. 2023. Evolutionary implications of C2photosynthesis: How complex biochemical trade-offs may limit C4evolution. Journal of Experimental Botany 74, 707-722.

Zhang YL, Zhang CL, Wang GL, Wang YX, Qi CH, Zhao Q, You CX, Li YY, Hao YJ. 2019. The R2R3 MYB transcription factor MdMYB30 modulates plant resistance against pathogens by regulating cuticular wax biosynthesis. BMC Plant Biology 19.

